# Multiple Geographical Origins of Environmental Sex Determination enhanced the diversification of Darwin’s Favourite Orchids

**DOI:** 10.1101/167098

**Authors:** Oscar Alejandro Pérez-Escobar, Guillaume Chomicki, Fabien L. Condamine, Jurriaan M. de Vos, Aline C. Martins, Eric C. Smidt, Bente Klitgård, Günter Gerlach, Jochen Heinrichs

## Abstract

Environmental sex determination (ESD) – a change in sexual function during an individual life span driven by environmental cues – is an exceedingly rare sexual system among angiosperms. Because ESD can directly affect reproduction success, it could influence diversification rate as compared with lineages that have alternative mating systems. Here we test this hypothesis using a solid phylogenetic framework of Neotropical Catasetinae, the angiosperm lineage richest in taxa with ESD. We assess whether gains of ESD are associated with higher diversification rates compared to lineages with alternative systems while considering additional traits known to positively affect diversification rates in orchids. We found that ESD has evolved asynchronously three times during the last ~5 Myr. Lineages with ESD have consistently higher diversification rates than related lineages with other sexual systems. Habitat fragmentation due to mega-wetlands extinction, and climate instability are suggested as the driving forces for ESD evolution.

## Introduction

Angiosperms have evolved a rich array of sexual systems that are unevenly distributed across lineages^1,2^. About 90% of angiosperms produce male and female reproductive organs in the same flower^2–4^. A small fraction of genera (i.e. 7%) include dioecious species, which bear sexes on separate individuals^5^. One unusual sexual system is referred to as labile sex expression, and involves changes in sex expression linked to environmental and phenotypic interactions^6^. A regular form of labile sex expression is the recurrent alternation of functionally male and female flower production over reproductive seasons^5^ (e.g. duodichogamy in *Acer*^6^). However, a more extreme form of labile sexual expression is environmental sex determination (ESD), where an individual can change its sex in response to environmental variables such as sunlight and temperature^3,6,7^. Therefore, plants with ESD can produce male, female or bisexual flowers either on the same individual or in separate individuals depending on specific environmental conditions^8^. ESD is a rare sexual system in angiosperms and has so far been reported for only *ca* 450 species in three families^5^. Several authors proposed that habitat heterogeneity afforded by fragmentation may provide conditions where ESD would be benefical^7–9^. Given the differential demand of resources by male (e.g. pollen production) and female (e.g. seed production) sex functions, ESD would enable the optimization of resource reallocation to either sex function in response to specific site characteristics.

Orchids, with about 25,000 species, are among the most species-rich families of flowering plants on earth^10–12^. They are well known for their diversity of reproductive systems^13^, but these remain poorly understood^14,15^. The vast majority of Orchidaceae produce male and female reproductive organs in the same flower^16^. Indeed, orchid flowers exhibit an extreme case of hermaphroditism in the plant kingdom because of the fusion of male and female organs into a gynostemium^17,18^. An exception to this rule however are the Catasetinae, a Neotropical orchid lineage containing 354 species^19,20^. Because of their striking sexual systems (i.e. ESD, protandry, adichogamy) and the diversity of pollination syndromes (e.g. male euglossine-bee pollination, oil bee pollination)^21,22^, Catasetinae have long attracted the attention of botanists and naturalists^23^. However, we still know little about the origin and evolution of ESD in this lineage.

Catasetinae includes eight genera, namely *Catasetum, Clowesia, Cyanaeorchis, Cycnoches, Dressleria, Galeandra, Grobya* and *Mormodes*, and most of the diversity is found in Central America and Amazonia regions^24^. Sex expression in Catasetinae, as in several other plant lineages with ESD, is regulated by ethylene production. The pioneer work of Gregg^25^ documented the role of sunlight exposure and ethylene synthesis on sex determination in *Cycnoches* and *Catasetum.* She convincingly demonstrated that ethylene concentrations are about two- to 100-fold higher in inflorescence apices that are fully exposed to sunlight compared with those kept in shade. Consequently, inflorescences with higher ethylene concentration usually produce female flowers, whereas those with lower levels produce male flowers^25^.

To date, about one third of the Catasetinae (i.e. *ca* 100 species) are known to have ESD. Therefore, the group provides a unique opportunity to study its evolutionary context in a phylogenetic framework. A recent study provided strong support for three independent origins of sexual plasticity in Catasetinae^14,26^. ESD evolved in *Catasetum, Cycnoches* and *Mormodes*, always from a protandrous ancestor. Protandry, in turn, was found to be the ancestral condition in the so-called core Catasetinae (*Catasetum, Clowesia, Cycnoches, Dressleria* and *Mormodes*). ESD could be a switch in the evolution of dioecy (i.e. spatial separation of sexes)^21^, as observed in other plant lineages with sexual plasticity such as *Fuchsia* and *Hebe*^27–29^.

It is still unclear whether the independent origins of ESD occurred synchronously and under the same environmental conditions in a single biogeographical region or evolved asynchronously in different regions, possibly under different environmental conditions. It also remains elusive whether lineages with ESD are associated with higher diversification rates compared with hermaphroditic lineages. In view of its rarity, we hypothesise that ESD could be a disadvantageous system leading to lower diversification rates. Alternatively, ESD could be evolutionarily unstable and could thus represent a transitional state rather than a stable reproductive system.

The extremely high orchid diversity is thought to be related with the evolution of novel traits and the invasion of novel environments^30^, possibly confounding analysis of the effects of ESD in orchid diversification. Therefore, we also investigated the impact of epiphytism^10,31^ and male euglossine-bee pollination^21,30^ on the diversification dynamics of Catasetinae. Epiphytism and male euglossine-bee pollination are traits that occur in all ESD species of Catasetinae, but these traits are shared with some Catasetinae species that have alternative reproductive systems^24^ (see *Methods*). Therefore, it is possible that these traits might have influenced the diversification of the subtribe, yet their specific contribution to diversification rates are still unknown.

The limited knowledge on the evolutionary dynamics of ESD in orchids stems partly from the lack of well-sampled species-level phylogenies^5^. This is particularly true for Catasetinae, in which previous studies did not attain even taxon sampling, thus leaving species-rich clades dramatically underrepresented (e.g. *Catasetum*; see^32–34^). In the current study, we generate the first densely sampled phylogeny of Catasetinae. We combine data on species distribution and a wide range of comparative phylogenetic methods to reconstruct the evolutionary history of ESD. We then infer the biogeographic history of Catasetinae, test whether their ESD lineages have higher diversification rates than non-ESD lineages, while accounting for the possible contribution of other traits to orchid diversification. We also investigate whether past and modern climate niche preferences of lineages with ESD are divergent from non-ESD lineages.

## Results

### *Phylogeny and molecular dating of* Catasetinae

Our DNA matrix included a total of 282 species, of which 132 belonged to Catasetinae (i.e. ~37% of the extant diversity of the subtribe; Tab. S1). Analyses based on Maximum Likelihood (ML) and Bayesian inference (BI) recovered similar topologies. However, they revealed several conflicting positions of species in the nuclear and plastid trees with bootstrap percentage values (MLBS) ≥ 75 and/or posterior probabilities (PP) ≥ 0.95 (Fig. S1-S2).Figure 1 and Figure S3 show the ML trees inferred from concatenated nuclear and non-conflicting plastid sequences, respectively. Here, after exclusion of conflicting positions from the DNA matrices (see *Methods* and Appendix S1 for a detailed explanation on incongruence handling), almost all backbone nodes of the phylogeny achieved MLBS ≥ 75 and PP > 0.95 support.

**Fig. 1.**
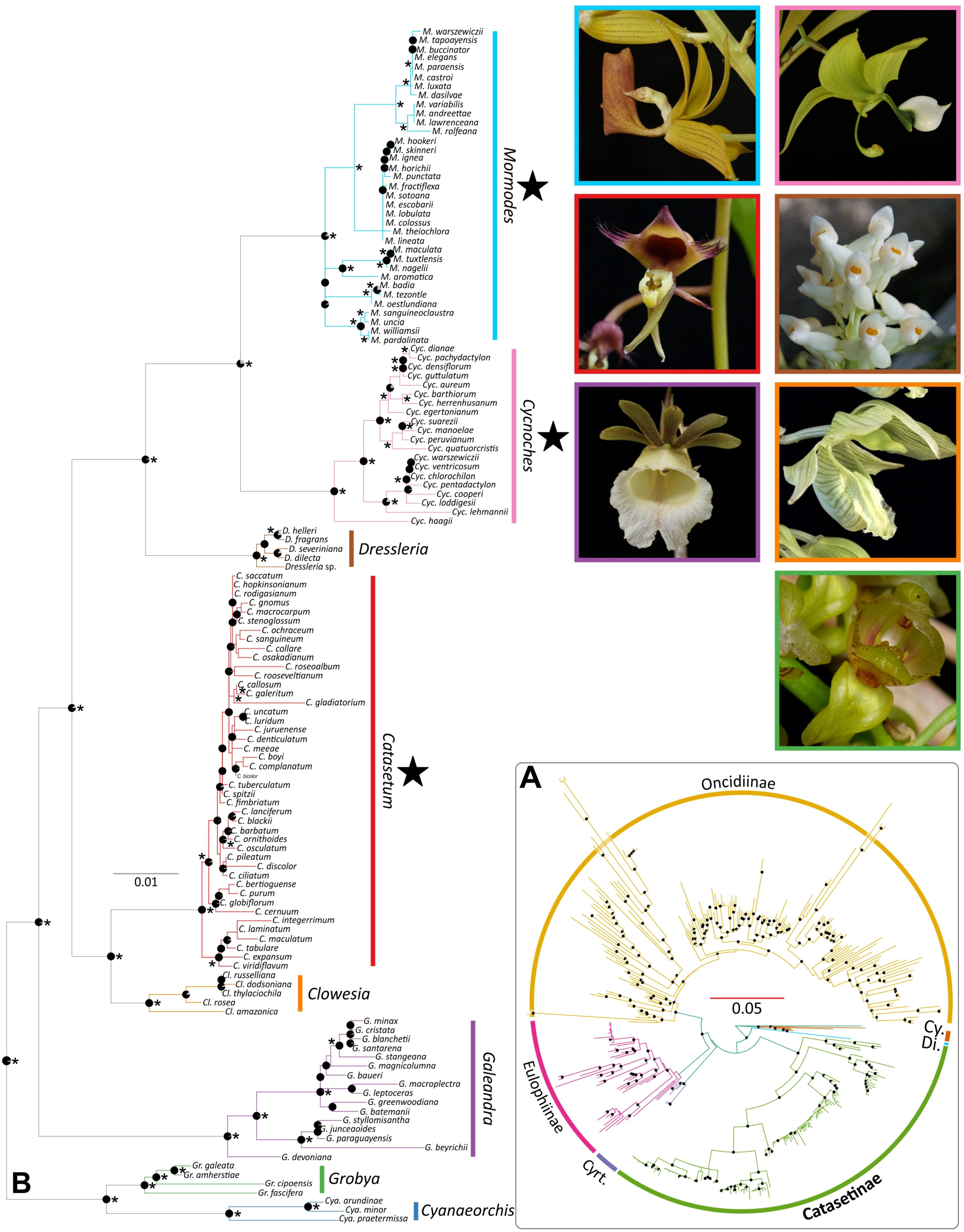
Best scoring ML tree of Catasetinae (B) and sister subtribes (i.e. Eulophiinae, Dipodiinae, Oncidiinae, Cyrtopodiinae, Cymbidiinae; A) obtained from non-conflicting concatenated nuclear and plastid loci. Node pie-diagrams indicate Bootstrap percentage values (MLBS > 75), where fully black diagrams indicate MLBS of 100. Asterisks represent BPP values higher than 0.95. Representatives of each clade in Catasetinae (except *Cyanaeorchis*) are shown in pictures [inset: *Catasetum: Ca. ochraceum; Clowesia: Cl. russelliana; Cycnoches: Cyc. lehmannii; Dressleria: D. eburnea; Galeandra: Ga. macroplectron; Grobya: G. galeata; Mormodes: M. punctata*]. Pictures by: O. Pérez and G. Gerlach.

Divergence time estimates performed under a Bayesian relaxed clock model yielded a mean Coefficient of Variation (CV) value for branch-specific substitution rates of 0.73, indicating substantial among-branch rate heterogeneity, confirming that the use of a relaxed molecular clock is appropriate. Mean ages and the associated 95% credibility intervals obtained under the relaxed clock model for all nodes are provided in Figure S4. The dating analyses showed that *(i)* the Catasetinae and Eulophiinae+Cyrtopodiinae shared a common ancestor in a time interval ranging from the early Miocene to the early Oligocene (25.93 Ma ± 5 Myrs), and *(ii)* the diversification of the Catasetinae most recent common ancestor (MRCA) started around 19.5 Ma ± 4 Myrs during the early Miocene.

### Geographical origin of ESD and biogeographical history of Catasetinae

Table S4 provides ML statistics for six biogeographical models as inferred using BIOGEOBEARS. The best fitting model was BayArea, including founder-event speciation (*J* parameter; LnL_best model_ = − 860.3 vs LnL_second best model_= −875.5; AIC_best model_ = 1727 vs AIC_second best model_ − 1757; Tab. S4). This model estimated south-eastern South America as the most likely ancestral area of Catasetinae (Fig. 2; Fig. S5). The same region was supported as the ancestral area of *Cyanaeorchis* and *Grobya.* In contrast, *Galeandra* and *Catasetum* inhabited larger areas between Amazonia+Guiana Shield+south-eastern South America and Amazonia+south-eastern South America, respectively. The origins of *Clowesia* and *Cycnoches* were estimated to be in Amazonia, whereas the remaining lineages of Catasetinae (i.e. *Dressleria, Mormodes*) originated in Central America (Fig. 2; Fig. S5).

**Fig. 2.**
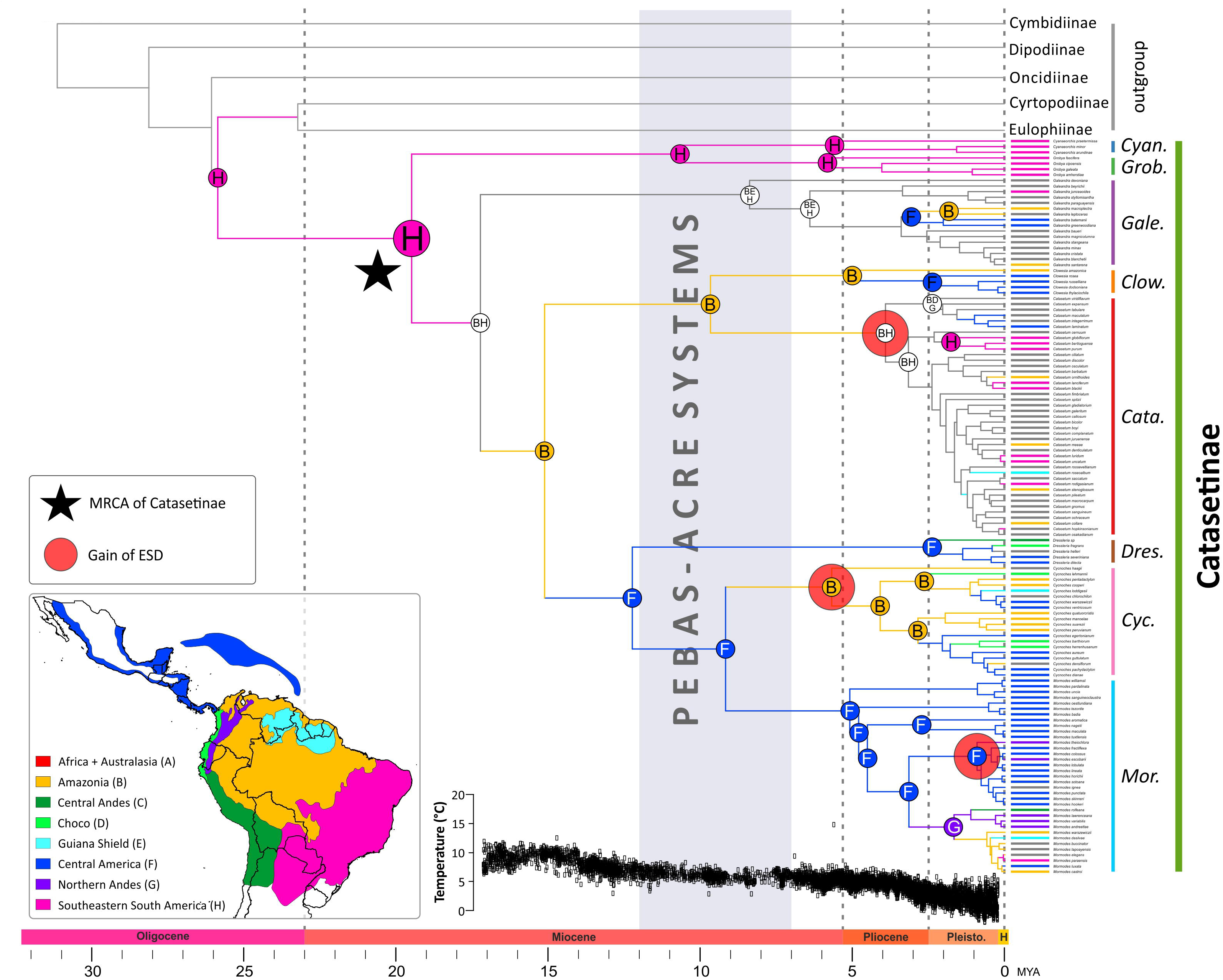
Chronogram of Catasetinae and sister subtribes obtained under a relaxed clock model, applied to a non-conflicting, concatenated nuclear and plastid loci. Minimum and maximum age intervals are provided in Fig. S4. Time scale is provided in million years (Mya). Node charts correspond to ancestral areas estimated under the BAYAREA+J model. The LCA of Catasetinae is indicated with a black star and gains of ESD are indicated with red circles. Coded biogeographical areas are colour-coded following inset map, and are shown in front of taxa names (grey colours indicate taxa distributed in more than one biogeographical region). The approximate time span of Pebas and Acre mega-wetlands^36^ is indicated by a grey rectangle. Inset: Coded areas used for biogeographical analysis. Geopolitical boundaries map generated by ArcMAP (http://www.sri.com) using political divisions and elevation data from DIVA-GIS (http://www.diva-gis.org/data). Climate curve of the last 17 Myr represented as a function of oxygen-isotope records^61^.

### Origin, climate niche and evolutionary dynamics of ESD

Ancestral character estimation analysis under the best model (i.e. single transition rate) identified three independent gains of environmental sex determination, without any losses (Fig. 2; Fig. S6). ESD first evolved during the early Pleistocene to late Miocene (5 Ma ±3 Myrs) at the MRCA of *Cycnoches* in the Amazonia region (Fig. 2; Fig. S5). Subsequently, two gains of ESD occurred towards the Pleistocene to Pliocene (3 Ma ±1 Myrs) and the Pleistocene (1 Ma ±1 Myrs) in Amazonia – south-eastern South America and Central America regions, respectively. These gains took place at the MRCAs of *Catasetum* and a small clade of *Mormodes* species restricted to Central America and Northern Andes, respectively (Fig. 2; Fig. S5, S6). Lineages with ESD further diversified mostly in lowland areas (i.e. Amazonia, Choco, south-eastern South America, Central America, Guiana Shield. Two episodic colonisations from Central America towards Northern Andes were recovered.

Bioclimatic variables for 1372 georeferenced occurrences of Catasetinae were adequately captured by the first two axes of a principal component analysis (PCA) (47.9% and 23.6%, respectively). The first PCA axis received contributions from all bioclimatic variables except BIO 3 (isothermality), BIO 4 (temperature seasonality), BIO 7 (temperature annual range), BIO 15 (precipitation seasonality), and altitude (Fig. S7). The second PCA axis mostly discriminated dry/hot environments, but also included contributions from BIO 15 (precipitation seasonality; Fig. S8). Therefore, discriminations between particular environments were not detected (Fig. S7-S9), as bioclimatic variables representing precipitation and temperature contributed similar amounts of variance to PCA axes 1 and 2 (**Fig. S7, S8**). Overall, the ordination of climatic niche space of lineages based on the bioclimatic variables indicated no evidence for general ecological segregation between lineages with and without ESD (**Fig. S9a**).

To further assess the apparent lack of climatic niche differentiation between ESD and non-ESD taxa, we excluded all auto-correlated variables (see *Methods*), and performed a non-metric dimensional analyses (NMDS) based on altitude, BIO 15 (precipitation seasonality), BIO 18 (precipitation of warmest quarter) and BIO 19 (precipitation of coldest quarter). Similarly, we found extensive overlaps between climatic niches of ESD and non-ESD lineages (Fig. S9b). However, climatic niche differentiation analysis using present-day bioclimatic variables are not directly informative of the climate niche preferences of ancestral lineages in which ESD arose. Thus, we performed ancestral character estimation analyses of twenty bioclimatic variables to assess whether ancestors having evolved ESD and those with alternative sexual systems diverged in their climatic niche preferences. We found that ancestral mean values of all bioclimatic variables were consistently similar between ancestors that independently evolved ESD and contemporary lineages with other sexual systems (Fig. 3). More importantly, ancestral mean bioclimatic values of lineages with and without ESD revealed little variation through time (Fig. S10).

**Fig. 3.**
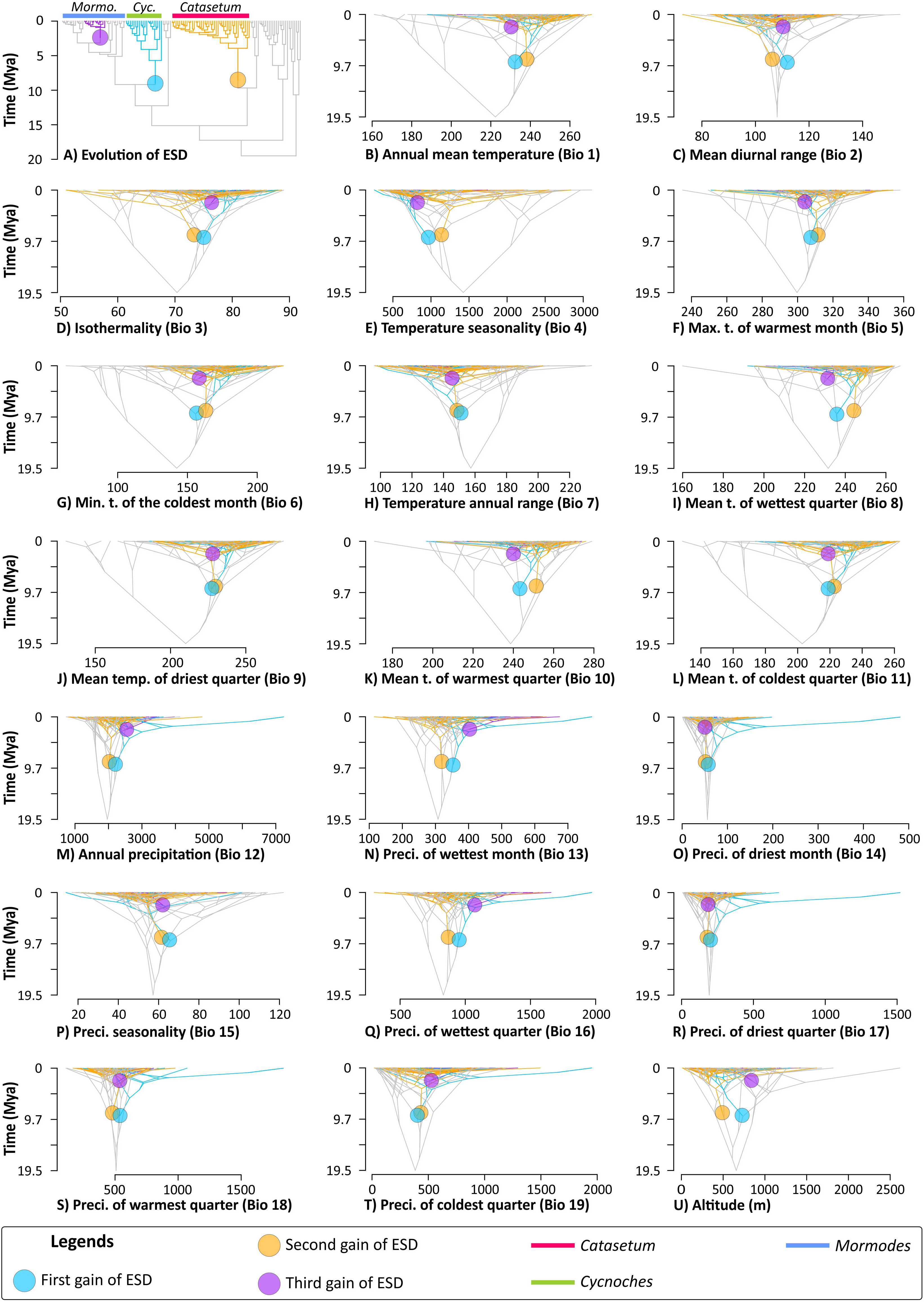
Evolution of ESD in Catasetinae and ancestral climatic-niche preferences depicted as traitgrams obtained via ACE analyses of mean values of bioclimatic variables. In all panels, coloured circles denote independent origins of ESD (colour-coded by chronological order). Colour coded branches denote lineages estimated to have ESD. Grey branches denote lineages inferred to have evolved alternative sexual systems (i.e. no ESD). Lineages in which ESD has evolved are color-coded accordingly. (A) Topological position of ESD origins (for posterior probabilities, see Fig. S6) (B) Annual mean temperature (Bio 1); (C) Mean diurnal range (Bio 2); (D) Isothermality (Bio 3); (E) Temperature seasonality (Bio 4); (F) Maximum temperature of warmest month (Bio 5); (G) Maximum temperature of coldest month (Bio 6); (H) Temperature annual range (Bio 7); (I) Mean temperature of wettest quarter (Bio 8); (J) Mean temperature of driest quarter (Bio 9); (K) Mean temperature of warmest quarter (Bio 10); (L) Mean temperature of coldest quarter (Bio 11); (M) Annual precipitation (Bio 12); (N) Precipitation of wettest month (Bio 13); (O) Precipitation of driest month (Bio 14); (P) Precipitation seasonality (Bio 15); (Q) Precipitation of wettest quarter (Bio 16); (R) Precipitation of driest quarter (Bio 17); (S) Precipitation of warmest quarter (Bio 18); (T) Precipitation of coldest quarter (Bio 19); (U) Altitude (meters). Note the little divergence through time of ancestral mean bioclimatic variables between lineages having ESD, and those with alternative sexual systems.

Trait-dependent diversification rate analysis supported a model with free speciation rates in lineages with and without ESD (LnL_best model_=−248.85; AICc_best model_=506.035 vs. LnL_null model_=−262.41; AICc_null model_=531.022, Tab. 1; Tab. S5). Under the best model, speciation rates were ~1.6-fold higher in lineages with ESD, while extinction rates were modelled under the same parameter (Fig. S11a; Tab. 1). We confirmed the robustness of these results through our tailored data simulations (see *Methods*), that generated an expectation about the distribution of ΔAIC values obtained from randomly reshuffled trait data (see *Extended Materials and Methods* of Appendix S1). The distribution of simulated ΔAIC did not overlap empirical AIC values. This suggests that our trait-dependent analysis is not biased by type I error (Fig. S14a).

**Table 1.**
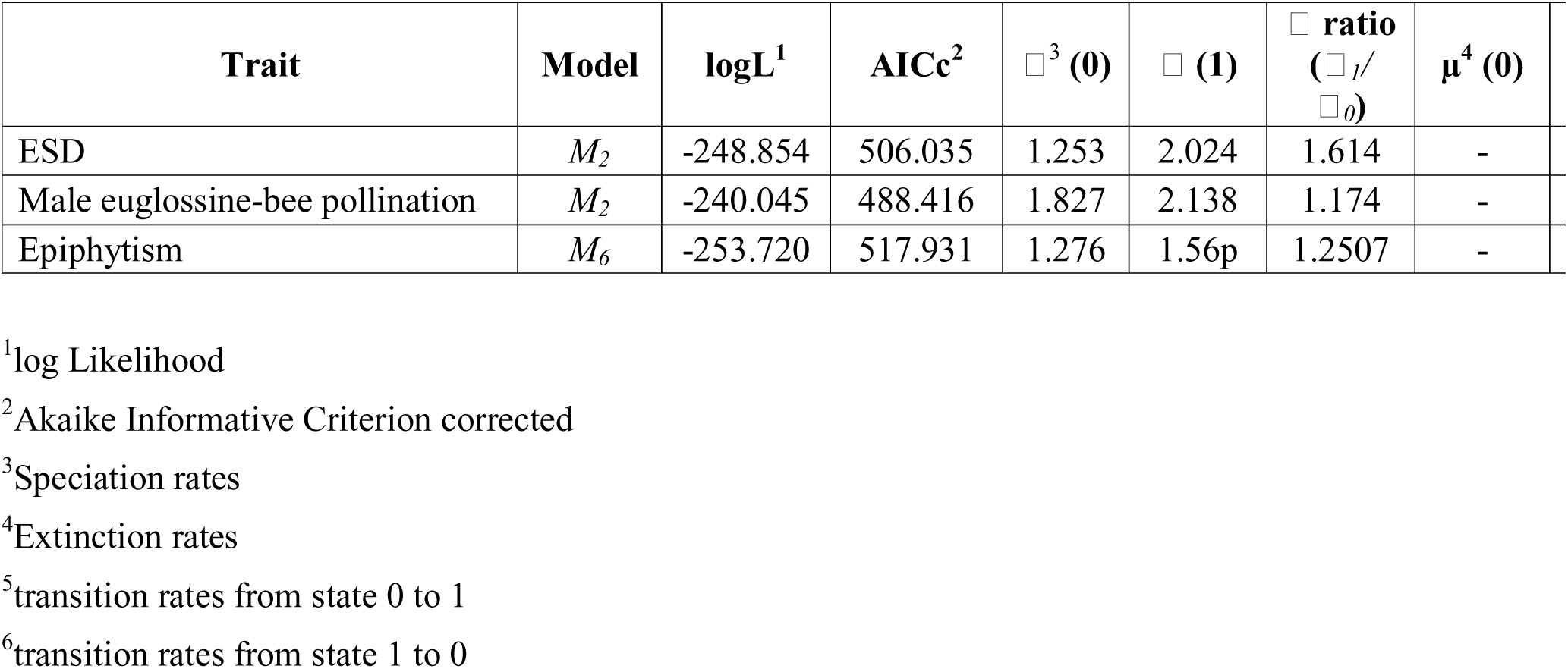
Best models and parameter values of BiSSE analysis on lineages with ESD vs. alternative sexual systems, epiphytism vs. other plant habits, and male euglossinebee pollination vs. other pollination systems.

### Evolutionary dynamics of male euglossine-bee pollination and epiphytism

Binary-state dependent diversification analyses of male euglossine-bee pollination and epiphytism supported models with free speciation rates and equal extinction and transition rates (*M_2_*; see *Methods*), and free speciation and transition rates but equal extinction rates (*M_6_*), respectively. Both models recovered slightly higher speciation rates for male euglossine-bee pollination (2.13 events/Myr; Fig. S12; Tab. 1; Tab. S6), and epiphytism (1.56 events/Myr; Fig. S13; Tab. 1; Tab. S7) than for lineages with alternative reproductive systems (1.82 events/Myr; Tab. S6) and plant habit (1.27 events/Myr; Tab. S7). Analysis of simulated data confirmed the robustness of real data analysis results for male euglossine-bee pollination. The distribution of ΔAIC values from real data and simulated data did not overlap: ΔAIC_simulated_ was centred towards 30 units while the ΔAIC_real data_ was estimated at 9 (Fig. S14b). Analysis of simulated data for epiphytism did not reach convergence, hence the veracity of binary-state dependent diversification analysis on the real data set could not be evaluated.

## Discussion

### Forest fragmentation as the context for the multiple geographical origins of ESD

Orchids have a rich variety of reproductive systems^35^, of which hermaphroditism is the most widespread in the family^16,35^, reflecting the angiosperm-wide pattern^2^. The Catasetinae, however, have evolved three reproductive systems (i.e. adichogamy, dichogamy, and ESD^14^), thus providing a unique opportunity to investigate biotic and abiotic factors shaping the evolution of these sexual systems. By relying on a solid phylogenetic framework, divergence time estimations, and ancestral character estimation, our study demonstrates the independent temporal and geographic origins of ESD in Catasetinae. More importantly, it reveals for the first time an apparent positive correlation between gains of sexual plasticity and higher speciation rates. However, our analyses show that other traits (i.e. male euglossine-bee pollination^21,30^, and perhaps epiphytism^30^), may also have positively contributed to the diversification of the Catasetinae.

Ancestral area estimation analyses suggest that sexual plasticity first originated in the Amazonian region during the late Miocene (5 Ma ±3 Myrs), a period preceded by the disappearance of large aquatic systems that dominated Amazonia from 17 to 7 Ma^36^. Such aquatic environments, known as Pebas Lake and Acre Systems, ranged from the drainage of western Amazonia to the Caribbean coast of Venezuela^37,38^ and prompted the fragmentation of pre-existing forests. The disappearance of Pebas and subsequent aquatic systems towards the late Miocene gave rise to flood-plains with a patchy vegetation dominated by palms, ferns and grasses^38,39^.

The late Miocene vegetation changes in the Amazonian region may also be related to the global cooling subsequent to the mid-Miocene Climatic Optimum (~15 Ma). These climate changes might have promoted forest fragmentation by range expansions and contractions of species due to dramatic temperature changes^37,40,41^. An example of the effect of drastic temperature changes on vegetation ranges is the peculiar *Campos Rupestres* in Brazil. There, dry periods provided the optimal conditions for vegetation to expand, allowing species to expand their distribution ranges, and subsequently occupy larger areas. Succeeding wetter periods would have prompted the contraction of distribution ranges, leading to the formation of isolated vegetation patches^42,43^. As a result, species like the orchids *Sarcoglottis caudata* and *Liparis beckeri* now occupy very restricted ranges, whereas other taxa have a disjunctive distribution, e.g. *Vellozia virgata*.

Sexual plasticity has been regarded as an adaption to patchy environments^7,8,44^, and this might have explanatory power for the gains of sexual plasticity in Catasetinae. The existence of patchy forests towards the end of the late Miocene in South America and vegetation changes in Central America might have facilitated the evolution of ESD by leading individuals to invest different amounts of resources to male and female gamete production. Resource reallocation to either function might thus have taken place as a function of an individual’s specific site characteristics^9,44^.

In a seminal paper, Zimmerman^45^ provided evidence towards the biased sex expression in *Catasetum viridiflavum.* In this species, female plants are more frequently distributed in open canopies whereas male plants often grow in closed canopies. Plants growing in open canopies are longer exposed to sunlight irradiation, thus favouring accumulation of energetic resources (i.e. plant biomass) that are later reallocated to female flower and fruit production^45^. The latter process requires considerable amounts of energy^46^ because of the massive number of seeds produced in a single capsule (see below). Sunlight irradiation appears to be the main ecological variable driving male/female flower production ratio in Catasetinae^25,45^. However, how other variables tied to resource constrains and patch quality (e.g. substrate type^45^, orchid-mychorriza interactions^47^) affect sex ratios in the Catasetinae deserves further research.

### ESD, epiphytism and male euglossine-bee pollination enhanced diversification in Catasetinae

The overwhelmingly low frequency of lineages with ESD compared to the vast majority of hermaphrodite angiosperms^5^ could suggest that sexual plasticity is a disadvantageous mating system or that it is evolutionary unstable. However, our study shows that Catasetinae clades with ESD have higher speciation rates than those with other sexual systems (i.e. adichogamy, protandry; Fig. S11). Here, the overarching comparison of speciation rates obtained from all analysed traits revealed a greater contribution of ESD evolution to the diversification dynamics of Catasetinae (rates ~1.6-fold higher than in lineages with other sexual systems). In contrast, the evolution of epiphytism (~1.2-fold higher) and male euglossine-bee pollination (~1.17-fold higher) affected diversification in the Catasetinae, but to a lesser extent.

Our ancestral character estimation analysis does not reveal any loss of ESD, suggesting that it is an evolutionary stable system in Catasetinae that has been maintained for up to 5 million years. Nevertheless, diversification in Catasetinae has also been prompted by the gain of traits that were earlier shown to influence diversification rates of Neotropical orchids in general (i.e. male euglossine-bee pollination^21^, epiphytism^30,31^). Taken together, these findings indicate that rate clade diversification is determined by several, potentially interacting factors^30^.

Vascular epiphytes are a key component of tropical forests^48^, and orchids account for 68% of the total vascular epiphyte diversity^49^. Epiphytism has evolved at least seven times since the Eocene in Orchidaceae^50^, and might have enhanced diversification by allowing colonization of new, largely unoccupied habitats such as branches and trunks of angiosperms^30^. In Catasetinae, epiphytism occurs in all members of the three most speciose genera, namely *Catasetum, Cycnoches* and *Mormodes*^24,51^ (290 species altogether20). It also occurs in all species of *Grobya*^24^ (five species), *Clowesia* (seven species) and six of the eleven species of *Galeandra*^52^. Epiphytic species from such clades inhabit in a wide array of ecosystems and biogeographical regions^24,53,54^. Contrastingly, terrestrial species are confined to species-poor clades with restricted distribution such as *Dressleria* (twelve species), *Cyanaeorchis* (three species), and a clade of five species nested in *Galeandra*^24,52,55^.

Pollination by male euglossine-bee evolved independently three times in Orchidaceae since the early Miocene^21^. It may have accelerated diversification rates by promoting switches in pollinators via small changes in the chemical profile of the floral scent, or alternatively by changes in pollinarium placement sites, hence allowing reproductive isolation^30^. Such pollination syndrome occurs in all species of *Catasetum, Clowesia, Cycnoches, Dressleria* and *Mormodes*, which altogether make up ~85% of total species number in Catasetinae. Alternative pollination syndromes (e.g. oil-bee pollination^22^, food deception) occur in the remaining, species-poor genera of Catasetinae (i.e. *Cyanaeorchis, Galeandra, Grobya*). Taken together, our results point to a prominent role of ESD in the evolutionary dynamics of the Catasetinae, evidenced by speciation rates ~1.6-fold higher than in orchids without ESD. Nonetheless, we do not exclude the additional contributions of traits, such as euglossine-bee pollination and epiphytism.

The reliability of our diversification rate results is strongly dependent on the power and hypotheses testing ability of our trait-dependant diversification model (i.e. BiSSE). The BiSSE method is known to be affected by within-clade pseudoreplication^56^, and sample size^57^, issues that are inherent to the number of independent origins of the analysed traits, and the sampling proportion of the clade under study, respectively. Thus, extreme care must be used when interpreting the outcome of trait-dependant diversification analyses^57^. However, by comparing different trait-diversification models with randomly reshuffled coding matrices in our simulations, and analysing other traits known to affect diversification rates in orchids, we have partially accounted for pseudo-replication bias. Indeed, our customized simulations revealed that BiSSE analyses of ESD and male euglossine-bee pollination are not affected by Type I errors. Nonetheless, diversification rate analyses can only account for the effects of the traits here considered, hence we cannot entirely rule out that other variables have additional effects^56^.

### ESD, protandry and adichogamy share the same climatic niche in Catasetinae

We have not detected significant differences in the ancestral and modern climate niches between Catasetinae lineages with ESD and other sexual systems (Fig. 3; Figs. S9-S10). Thus, particular temperature or precipitation regimes do not appear to have favoured any of these lineages. Species with ESD that occur often in habitats with longer sunlight exposure compared to species without ESD, produce bigger pseudobulbs, female flowers and fruits^45,58^. These species also produce larger capsules than Catasetinae lineages with protandry or adichogamy (e.g. *Cyanaeorchis, Clowesia, Dressleria*; Pérez-Escobar, pers.obs.). For instance, the capsules of *Cycnoches chlorochilon* produce on average up to three times more seeds (3,770,000) than fruits of *Cymbidium tracyanum* (850,000), an adichogamous species closely related to Catasetinae^46^. Therefore, under the same climatic niche, Catasetinae lineages with ESD might have produced more offspring than lineages with other sexual systems. However, additional data on fruit size and particularly on seed content for all Catasetinae members are needed to statistically test such assumption.

Occurrence of ESD requires a heterogeneous habitat (e.g. fragmented forest matrix), because under homogenous ecological conditions (e.g. forest understory) plants with ESD will bear predominantly one sex form, and mating success may thus be dramatically reduced. This highlights the important role of climate instability and its effects on habitat heterogeneity since the Miocene Climate Optimum in the evolution and maintenance of ESD. Nevertheless, it is likely that bioclimatic variables do not represent relevant environmental aspects of the niche in which ESD orchids occurs as compared with non-ESD lineages. Also, the climatic niche analyses pinpoint the explanatory power of variables with maximum variance, hence they will be mostly represented in PCA axes 1 and 2. However, such variables are not necessarily the most relevant and informative for our group of study. Therefore, even though ESD and non-ESD species niches largely overlap, ESD lineages might still be dependent on more specific ecological variables that are not necessarily captured by bioclimatic variables but that may yet be driven by climate (e.g. those operating on forest edge matrices^59^, or affecting physiognomic properties of vegetation).

## Conclusions

Environmental Sex Determination (ESD) in Catasetinae has long attracted and even confused^60^ botanists and naturalists23, and the evolutionary and biogeographic context for the evolution of this remarkable trait remained elusive before our study. Our results show that the evolution of ESD involved multiple gains in three biogeographical regions between late Miocene and late Pleistocene - geological periods that were marked by the extinction of pivotal aquatic environments and mega-wetlands^36,39^ and dramatic fluctuations in temperature^61^. Fragmentation of vegetation prompted by the demise of these aquatic systems in South America and by dramatic changes in temperature across the Neotropics might have fostered the evolution of sexual plasticity in Catasetinae. In spite of the extreme rarity of sexual plasticity among the rich array of angiosperm reproductive systems^5^, evolution of ESD in Catasetinae appears to be correlated with higher speciation rates. Other traits such as male euglossine-bee pollination and epiphytism, however, also appear to have positively influenced the spectacular diversity of Catasetinae. This illustrates the difficulty in isolating the effect of one trait on diversification rates from effects of others. The genetic contribution of an individual to future generations is enhanced by ESD because it promotes the development of the sex most suited for specific environmental conditions (i.e. sunlight intensity in Catasetinae^8,44^). We demonstrate that ESD-bearing Catasetinae inhabit the same climatic niche as their non-ESD relatives. Thus, plants with ESD might produce more offspring over plants having alternative sexual systems because of the increased opportunities for optimizing resource allocation to male and female function at each growing site.

## Materials and methods

### Taxon sampling, DNA sequencing and phylogenetic analyses

Species names, geographical origins, voucher specimens, and GenBank accession numbers of sequences included in phylogenetic analyses are provided in Table S1. We built our alignment based on the molecular datasets of Whitten et al.^33^, Salazar et al.^51^, Neubig et al.^62^, Bone et al.^63^, and Pérez-Escobar et al.^34,64^, and further generated 57 nuclear ribosomal and 25 plastid sequences for poorly sampled lineages such as *Catasetum, Dressleria* and *Grobya* (Tab. S1). Because the precise position of Catasetinae inside the tribe Cymbidieae was elusive^33,55^, we also sampled selected representatives of closely related subtribes (i.e. Cyrtopodiinae, Dipodiinae, Eulophiinae, and Oncidiinae). Genomic DNA extraction, amplification, sequencing and alignment procedures are the same as in Irimia et al.^65^, Pérez-Escobar et al.^14^, and Bechteler et al.^66^. For the newly sampled taxa, we sequenced nuclear ribosomal external and internal transcribed spacers (ETS and ITS, respectively), a fragment of the *Xdh* gene, and also a ~1500 bp fragment of the plastid gene *ycf*1, and the *trn*S–*trn*G intergenic spacer. Amplification settings and sequencing primers used for ITS, ETS, *Xdh, trn*S– *trn*G, and *ycf*1 are specified in Table S2.

Loci were aligned using MAFFT 7.1^67^ and were further manually checked, yielding a total matrix of 281 taxa and 8104 nucleotides (~24% parsimony informative sites). Congruence between nuclear and plastid datasets was assessed following Pérez-Escobar et al.^34^, and using PACo^68^ (Appendix S1). The procedure is available as a pipeline (http://www.uv.es/cophylpaco/) and was also employed to identify operational terminal units (OTUs) from the plastid dataset conflicting with the nuclear dataset (potential outliers detected by PACo are shown in Fig. S1-S2). A detailed explanation on PACo and a rationale on outlier handling is provided in the Extended Materials and Methods section of Appendix S1.

Phylogenetic analyses of separate and concatenated loci were carried out under maximum likelihood (ML) and Bayesian inference (BI), following Pérez-Escobar et al.^14,34^, and Feldberg et al.^69^. The best-fitting substitution models for ML and Bayesian analyses were obtained for each data partition using jModelTest v.2.1.6^70^ (Tab. S3). Phylogenetic inference relied on the ML and BI approaches implemented in RAxML-HPC 8.2.4^71^ and MrBayes 3.2.272, respectively, and were carried out on the CIPRES Science Gateway computing facility^73^. Maximum likelihood bootstrap supports (MLBS) were generated for the ML tree and MLBS ≥ 75% considered as good support, and Bayesian Posterior Probabilities (PP)≥ 0.95 for the Bayesian majority rule consensus topology were regarded as significant^74,75^.

### Molecular clock dating

Divergence time estimates were conducted using the Bayesian relaxed clock approach of BEAST 2.1.3^76^ with a concatenated nuclear+plastid subset of the data obtained after the PACo analysis. Strict and uncorrelated lognormal molecular clock models, both with pure-birth speciation models as recommended for species level sampling^77^, were compared to explore clock-likeness of the data. For calibrating the relaxed clock model, there are few fossils unambiguously assigned to Orchidaceae^78^, and these are placed to lineages very distantly related to Catasetinae (i.e. *Dendrobium, Earina*, both Vandeae). Recently, a new putative orchid fossil from the Dominican amber (early - middle Miocene, 15-20 Ma) was described and assigned to the tribe Cymbidieae^79^. However, the author provided no evidence to unambiguously assign such fossil to Cymbidieae or orchids in general. Secondary calibrations are therefore the only option for dating the tree of Catasetinae. To this end, we relied on the age estimates obtained from a fossil-calibrated global phylogeny of Orchidaceae^50^, which were set with normally distributed priors that reflected uncertainty in the primary analysis^76^. Settings for absolute age estimation are detailed in the *Extended Materials and Methods* of Appendix S1.

### Ancestral areas estimation

Species ranges were coded from the literature^19,60^ and from herbarium specimens housed in the herbaria AMES, COL, F, M, MO, SEL, US, using the R-package SpeciesGeoCoder^80^. Distribution data were also obtained from own field observations. Biogeographical areas were defined considering putative areas of endemism of Catasetinae as well as species distributions observed in other plant lineages such as Rubiaceae^37^ and Bromeliaceae^81^. We divided the geographical range of Catasetinae and outgroup taxa in eight biogeographic regions (see *Extended Materials and Methods* of Appendix S1 for a detailed description of coded areas): (1) Central America; (2) Guiana Shield; (3) Amazonia^37^; (4) Chocó; (5) Northern Andes; (6) Central Andes; (7) South-eastern South America; and (8) Africa and Australasia. We used the R-package BIOGEOBEARS (Biogeography with Bayesian and Likelihood Evolutionary Analysis in R script^82^) to estimate ancestral areas on the phylogeny of Catasetinae.

### Climatic niche analyses

We mapped georeferenced collection records obtained from floras, GBIF database and herbarium specimens (mean of five specimens per species, maximum number of record per species set to ten), and they represent the known distribution of extant Catasetinae species included in our taxon sampling. To query GBIF database, we relied on the function *occ* of the R-package SPOCC^83^. We further extracted corresponding values of altitude and 19 climatic variables (30 seconds resolution) reflecting temperature and precipitation regimes from the WorldClim database (available at: http://www.worldclim.org/current), using the function *extract* of the R-package RASTER84. To characterise the climate-niche occupied by Catasetinae lineages with ESD or other sexual systems, we performed Principal Component Analysis (PCA) on the mean values of the 20 environmental variables of each species, using the default function *princomp* of the R (Fig. S7). We further explored the contribution of bioclimatic variables to axes with the most variance by plotting the loadings of every bioclimatic variable onto the axes (Fig. S8, S9).

To avoid spurious results arising from inclusion of correlated variables 63,85, we determined the Pearson’s correlation coefficients between the variables and altitude and then included only variables with a Pearson’s correlation coefficient <0.5, taking a single variable in correlated clusters. This way, we selected altitude, BIO 15 (precipitation seasonality), BIO 18 (precipitation of warmest quarter) and BIO 19 (precipitation of coldest quarter) as a set of maximally uncorrelated variables. We analysed these variables using the R-package VEGAN^86^ to perform non-dimensional metric scaling analyses (NMDS) using the dataset of 1372 georeferenced herbarium specimens.

### Ancestral character estimation

We coded for absence (state 0) and presence (state 1) of ESD in 131 species of Catasetinae, plus eight taxa of Cyrtopodiinae (see below). We sampled 68 of the 164 known species with ESD in Catasetinae (i.e. ~41% of the ESD diversity). Information on occurrence of ESD was obtained from the literature^24,58,87^. We estimated the origin of ESD within Catasetinae using a stochastic character mapping approach implemented on a maximum likelihood framework for ancestral character estimation analysis (ACE) implemented by the function *make.simmap* in the R-package PHYTOOLS88. Under this approach, we fitted single (ER), symmetrical (SYM), and asymmetrical character transition rate models (ARD) and performed stochastic mapping on 1000 iterations using the maximum clade credibility tree derived from the BEAST dating analysis (see above). Eight members of the Neotropical genus *Cyrtopodium* were selected as outgroup taxa, and they exhibit hermaphroditic flowers (i.e. no ESD).

To assess whether ancestors with ESD and those with alternative sexual systems differed in their climatic niche preferences back in time, we performed ACE analyses of the same nineteen bioclimatic variables plus altitude, obtained from georeferenced occurrences (see *Climate niche analyses* section of *Methods*). ACE of absolute mean values relied on a Maximum Likelihood framework, and were performed on a phylogeny derived from the character stochastic mapping analysis of ESD (using ER as transition rate model, see *Results*). We then visualized the variation through time of all variables (**Fig. 3**), using the function *phenogram* of the R-package PHYTOOLS88. We also plotted mean ancestral values and their corresponding 95% confidence intervals (CI) estimated for every node of the Catasetinae phylogeny, using the R-package GGPLOT2^89^ (Fig. S10).

### Trait-dependent diversification analyses

We tested whether linages with ESD (state 1) had higher speciation rates than hermaphroditic, monoecious clades (state 0). In addition, to tease apart the contribution of biotic variables to Catasetinae diversification that are known to positively influence diversification rates in orchids, we investigated the evolutionary dynamics of male euglossine-bee pollination and epiphytism in the subtribe. The former trait occurs in all species of the genera *Catasetum, Clowesia, Cycnoches, Mormodes* and *Dressleria*^21,24,90^, while the latter occur in all species of *Catasetum, Clowesia, Cycnoches, Mormodes, Dressleria*^24^, and part of *Galeandra* species52. We relied on the Binary State Speciation and Extinction (BiSSE) model^91^ to estimate diversification rates associated with each trait. We used the R-package DIVERSITREE^91^ to perform eight diversification models in the maximum likelihood framework on 100 randomly sampled dated trees from the Bayesian divergence times analysis. We tested eight models with different configurations of speciation, extinction, and transition rates between characters. Detailed settings for binary-trait dependent diversification analyses, and simulations to account for type I error biases^92^ are provided in Appendix S1.

## Acknowledgements

We are grateful to Martina Silber (Munich, Germany) for valuable assistance provided during lab work, to Mario Blanco (San José, Costa Rica), Diego Bogarín (San José, Costa Rica), Norman Cash and family (Managua, Nicaragua) for assistance during field trips, and to Gustavo Romero (Herbarium HUH, Cambridge, U.S.A) and Mario Blanco (University of Costa Rica) for providing plant material. Adarilda Petini-Bennelli provided vegetal material of Catasetinae. Paula Rudall and Richard Bateman critically read the manuscript. To R. Camara, for discussions. The handling editor and two anonymous reviewers for providing critical comments to the manuscript. This work was supported by the Colombian Administrative Department of Science and Technology (COLCIENCIAS; granted to OAPE, ref. 512). F.L.C. has benefited from an “Investissements d’Avenir” grant managed by Agence Nationale de la Recherche (CEBA, ref. ANR-10-LABX-25-01).

## Author contributions

O.A.P.E., G.C., F.L.C., J.V. and J.H. designed research; A.M and E.S. collected samples; A.M., E.S. and O.P. performed the lab work; O.A.P.E., F.L.C., and G.C. performed all analyses; O.A.P.E., G.C., F.L.C., J.V., B.K., J.H., G.G., A.M., and E.S. wrote the manuscript.

## Supplementary material

**Appendix S1.** Extended Materials and Methods, supplementary tables and figures.

